# The DBL-1/TGF-β signaling pathway regulates pathogen-specific innate immune responses in *C. elegans*

**DOI:** 10.1101/2021.03.30.437693

**Authors:** Bhoomi Madhu, Tina L. Gumienny

## Abstract

Innate immunity in animals is orchestrated by multiple cell signaling pathways, including the TGF-β superfamily pathway. While the role of TGF-β signaling in innate immunity has been clearly identified, the requirement for this pathway in generating specific, robust responses to different bacterial challenges has not been characterized. Here, we address the role of DBL-1/TGF-β in regulating signature host defense responses to a wide range of bacteria in *C. elegans*. This work reveals a role of DBL-1/TGF-β in animal survival, organismal behaviors, and molecular responses in different environments. Additionally, we identify a novel role for SMA-4/Smad that suggests both DBL-1/TGF-β-dependent and -independent functions in host avoidance responses. RNA-seq analyses and immunity reporter studies indicate DBL-1/TGF-β differentially regulates target gene expression upon exposure to different bacteria. Furthermore, the DBL-1/TGF-β pathway is itself differentially affected by the bacteria exposure. Collectively, these findings demonstrate bacteria-specific host immune responses regulated by the DBL-1/TGF-β signaling pathway.

## INTRODUCTION

Living organisms recognize and respond to potential environmental insults by coordinating protective defenses^1^. Invertebrates and vertebrates both employ conserved innate immune response as immediate front-line protection from challenges including pathogenic bacteria^2–5^. These responses are tailored to the bacterial challenge. However, how these responses are specified and what the responses are to different pathogens remains a challenge^6^.

The roundworm *Caenorhabditis elegans* is an established model system to study regulation of immune responses in vivo^7^. *C. elegans* naturally thrives in a soil environment where it feeds on bacteria and is in constant association with a diverse range of microbes that are both food and threat^3,7^. A limited number of pathogens are known to infect *C. elegans*, including mycobacterial, Gram-negative, and Gram-positive bacterial species, and fungi^8–15^. *C. elegans* has an innate immune system that confers protection through behavioral, physical (exoskeleton), and molecular mechanisms^7^.

Infection in *C. elegans* induces molecular immune defenses coordinated by conserved Toll-like receptors, MAPK (mitogen-activated protein kinase) signaling, insulin-like signaling, and DBL-1/TGF-β (transforming growth factor β) signaling^8–13,16,17^. These pathways regulate an overlapping set of target defense genes, indicative of coordinated crosstalk between these signaling pathways. While the roles of Toll, MAPK, and insulin-like signaling pathways in immune responses to a wide variety of bacteria are well characterized in *C. elegans* and other organisms, a role for DBL-1/TGF-β signaling in eliciting robust targeted immune responses to different bacterial challenges has been identified but is not well defined^16,18–22^. Previous reports indicate that overexpression of *dbl-1* enriches expression of many immune response genes including lectins, saposin-like proteins, and lysozymes^11,23–25^. While the role of DBL-1 in defending nematodes from a few Gram-negative bacteria is reported, its possible role in protection against Gram-positive bacterial infection has not been well characterized^20,26,27^.

In this work, we examined the role of DBL-1/TGF-β signaling in regulating an array of microbe-specific immune responses. Using behavioral and molecular approaches, we identified DBL-1-dependent and -independent immune responses that are tailored to the specific bacterial exposure. We also identified a non-canonical role for the DBL-1 pathway transcription regulator SMA-4 in an avoidance response to specific bacteria. Additionally, we show that DBL-1 signaling is induced in response to Gram-negative bacteria but is repressed in response to Gram-positive bacteria. We propose that animals lacking DBL-1 signaling respond with heightened avoidance behaviors to selected bacterial environments because they perceive the environment as more hostile. Collectively, our findings highlight a central role for DBL-1 in regulating a suite of bacteria-specific host defenses and also demonstrate bacteria-responsive regulation of DBL-1 signaling.

## MATERIALS AND METHODS

### Strains and maintenance

#### *C. elegans* strains

All *C. elegans* strains were maintained on EZ media plates at 20°C^28^. *C. elegans* strains were maintained without contamination and starvation for at least five generations before every experiment. Supplementary Table 1 includes the list of all strains used in this study. These strains were generated by standard genetic crosses and confirmed by small body size phenotype and presence of fluorescence.

#### Bacterial strains

The bacterial strains used in this study include *Bacillus megaterium* (Carolina Biological Supply Company), *Escherichia coli* (OP50), *Enterobacter cloacae* (49141TM), *Enterococcus faecalis* (51299TM), *Klebsiella oxytoca* (49131TM), *Serratia marcescens* (Carolina Biological Supply Company), and *Staphylococcus epidermidis* (49134TM). *E. faecalis* in brain heart infusion media and all other bacteria in tryptic soy broth were grown for nine hours at 37°C as previously described^28^. Bacterial cells were pelleted at 5000 rpm for 15 minutes and concentrated twenty-fold. EZ media plates were freshly seeded with concentrated bacteria in full lawns. The plates were incubated at 37°C overnight before using for experiments.

### Lifespan assay

Lifespan assay was performed as previously described^29–31^. Concentrated bacterial cultures were spread on 6 cm diameter EZ media plates (full lawn plates) containing 50 μg/ml 5-fluorodeoxyuridine (FUdR) to cover the surface of the plates entirely. Wild-type and *dbl-1(−)* animals (n= at least 30) were fed on control and test bacteria on full lawn plates at the L4 stage in quadruplicate. The plates were scored for live and dead nematodes every 24 hours until all animals were dead. Animals were scored as dead if they did not respond to gentle touch with a sterilized platinum wire and were removed from the plate. At least three independent trials were performed. Worms that died by desiccating on the walls of the plates were censored from the analysis.

### Pharyngeal pumping rate

Wild-type and *dbl-1(−)* L4 animals (n=12) were fed on control and test bacteria on full lawn plates. The number of contractions of the pharyngeal bulb was counted for 20 seconds to calculate the pharyngeal pumping rate of animals. Two counts were made and averaged for each animal. Three independent trials were performed in triplicate^32^.

### Intestinal barrier function assay

The intestinal barrier function assay was performed as previously described^33^. Wild-type and *dbl-1(−)* L4 animals were fed on control and test bacteria on full lawn plates. The assay was performed when about 50% of the population with the lowest mean lifespan remained alive. At least fifteen animals were sampled at the specified times to examine intestinal tissue integrity. The intestinal barrier integrity was assessed using a blue dye, eriogluacine disodium salt (5% wt/v), as an indicator of tissue integrity as the animals age. Leaking of this blue dye outside the intestinal lumen indicates reduced intestinal integrity. The animals were washed with S buffer and were incubated in erioglaucine disodium salt solution in a 1:1 ratio for 3 hours. The animals were then washed thrice with S buffer and were mounted on 2% agarose pads on glass slides. 10 μM levamisole was added to paralyze the animals. The animals were imaged on a Nikon DS-Ri2 camera mounted to a Nikon SMZ18 dissecting microscope. The leakiness of the intestine was assessed and scored as ‘1’ for no leakage/no Smurf, ‘2’ for mild leakage/mild Smurf, and ‘3’ for severe leakage/severe Smurf phenotypes. The experiment was performed in at least three independent trials for each experimental condition.

### Microbial avoidance assay

Microbial avoidance assays were performed as previously described^34^. 20 μl of the concentrated bacterial cultures were spotted on 6 cm diameter EZ media plates and incubated at 37°C overnight. Wild-type, *dbl-1(−)*, *sma-2(−)*, *sma-3(−)*, and *sma-4(−)* L4 hermaphrodites were placed on control and test bacterial lawn (n=30 per condition/trial, performed in triplicate). The plates were scored for number of worms occupying the lawn at the indicated time points. Three independent trials were performed. The avoidance ratio (A) was calculated using the formula:

A= number of animals off the lawn/ total number of animals Avoidance of populations was scored as mild if less than 40% of the population was not on the food source (A<0.4), as moderate avoidance if 0.4<A<0.6, and as strong avoidance if A>0.6.

### RNA isolation

Animals were synchronized as embryos by bleaching mixed-stage populations^35^. Total RNA was extracted from animals at 48 hours after the L4 stage. Total RNA was extracted by the freeze cracking method as previously described^36^.

### Differential expression analysis by RNA sequencing

RNA from wild-type and *dbl-1(−)* adult populations fed on control and test bacteria was extracted in three independent trials. Sequencing libraries from the extracted RNA were generated using the NEBNext^®^ RNA Library Prep Kit for Illumina^®^ (NEB, USA) following manufacturer’s recommendations. 1 μg RNA of each sample was used as input material for the RNA sample preparations. Novogene performed RNA sequencing of samples. Differential expression analysis of wild-type compared to *dbl-1(−)* populations grown on different bacteria was performed using the DESeq R package (1.18.0)^37^. Genes with an adjusted *p*-value < 0.05 found by DESeq were assigned as differentially expressed.

### cDNA synthesis and qRT-PCR

After RNA isolation, cDNA was synthesized and quantitative real-time PCR was performed as previously described^38^. 2 μg of total RNA isolated was primed with oligo(dT) and reverse transcribed to yield cDNA using the SuperScript III reverse transcriptase kit as per manufacturer’s protocol (Invitrogen). Real-time PCR was performed on a QuantStudio3 system (Applied Biosystems by Thermo Fisher Scientific) using the PowerUP SYBR Green master mix (Applied Biosystems) according to manufacturer’s instructions. Three independent biological trials were performed. Each biological trial was performed in three technical replicates for each condition. Primer sequences are available in Supplementary Table 2. QuantStudio Design and Analysis Software v1.5.1 was used to calculate raw C_t_ values. The C_t_ values for the target genes were normalized to the housekeeping gene *act-1* (actin) (Applied Biosystems by Thermo Fisher Scientific). Fold change in gene expression between experimental sample and the control was determined by using the formula: 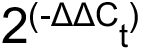.

### Imaging

RAD-SMAD reporter strains were placed on full lawns of the control or test bacteria at the L4 stage. L2 progeny were mounted on 2% agarose pads and anesthetized by using 1 mM levamisole and fluorescence was captured by a Zeiss LSM 900 confocal microscope using a 40X oil objective. At least fifteen animals with at least five hypodermal nuclei per worm in the focal plane were imaged per condition, giving a moderate effect size as determined by power analysis. The experiment was performed in three independent trials. The microscope conditions were optimized with respect to the control and test conditions and kept consistent within each trial. Mean fluorescence intensities were measured as previously described using the Zeiss ZEN lite software^39^.

The innate immune reporter strains were transferred to full lawns of the control and test bacteria at the L4 stage and were imaged after 48 hours of exposure. Fluorescence of the reporter strains was captured by a Nikon DS-Ri2 camera mounted on a Nikon SMZ18 dissecting microscope. Animals were mounted on 2% agarose pads and anesthetized with 1 mM levamisole. At least fifteen animals were imaged per condition as determined by power analysis with a moderate effect size. The microscope conditions were optimized with respect to the control and test conditions and kept consistent within each trial. However, imaging exposure times were different between some trials to prevent saturation of signal in experimental conditions. Three independent trials were performed. Mean fluorescence intensities were measured using the Nikon NIS Elements AR v5.02 software.

### Statistical analyses

Lifespans of *C. elegans* populations were calculated by the Kaplan-Meier method and statistical analysis was performed using the two-tailed log-rank test. The average pharyngeal pumping rates were compared by the two-tailed unpaired *t*-test. The intestinal barrier function phenotypes were statistically analyzed by the Chi-square test. The avoidance ratio was compared by repeated measures ANOVA using Tukey’s post-hoc test. qRT-PCR values and mean fluorescence intensities were evaluated using the two-tailed unpaired *t*-test. RNA sequencing analysis was performed using the DESeq R package (1.18.0). The resulting *p*-values were adjusted using the Benjamini and Hochberg’s approach for controlling the false discovery rate.

## RESULTS

### Loss of DBL-1 reduces lifespan of animals fed on specific bacteria

To study the requirement for DBL-1 in specific responses to pathogens at behavioral, molecular, and physiological levels, we first established a panel of bacteria for innate immune studies in *C. elegans* that would facilitate genetic and molecular studies over time: previous studies have been limited in range of challenge and used a pathogen that killed animals in hours or a few days. The control bacteria chosen was Gram-negative *E. coli* OP50, a commonly used strain for laboratory culture of *C. elegans*. The panel of test bacteria comprises Gram-negative and -positive bacteria that are opportunistic pathogens in humans and are found in the natural habitat of *C. elegans*^40^. We selected three Gram-negative test strains, *S. marcescens*, *E. cloacae*, and *K. oxytoca*, and three Gram-positive test strains, *B. megaterium* and *E. faecalis*, and *S. epidermidis*.

We first asked if our panel of opportunistic bacteria affect lifespan of *C. elegans*. Wild-type animals on Gram-negative *S. marcescens* have lifespans comparable to *E. coli*-fed animals. Interestingly, we noted an extended lifespan of wild-type animals on the other two Gram-negative strains and all three Gram-positive strains (Fig.1, Supplementary Table 3).

**Figure 1.**
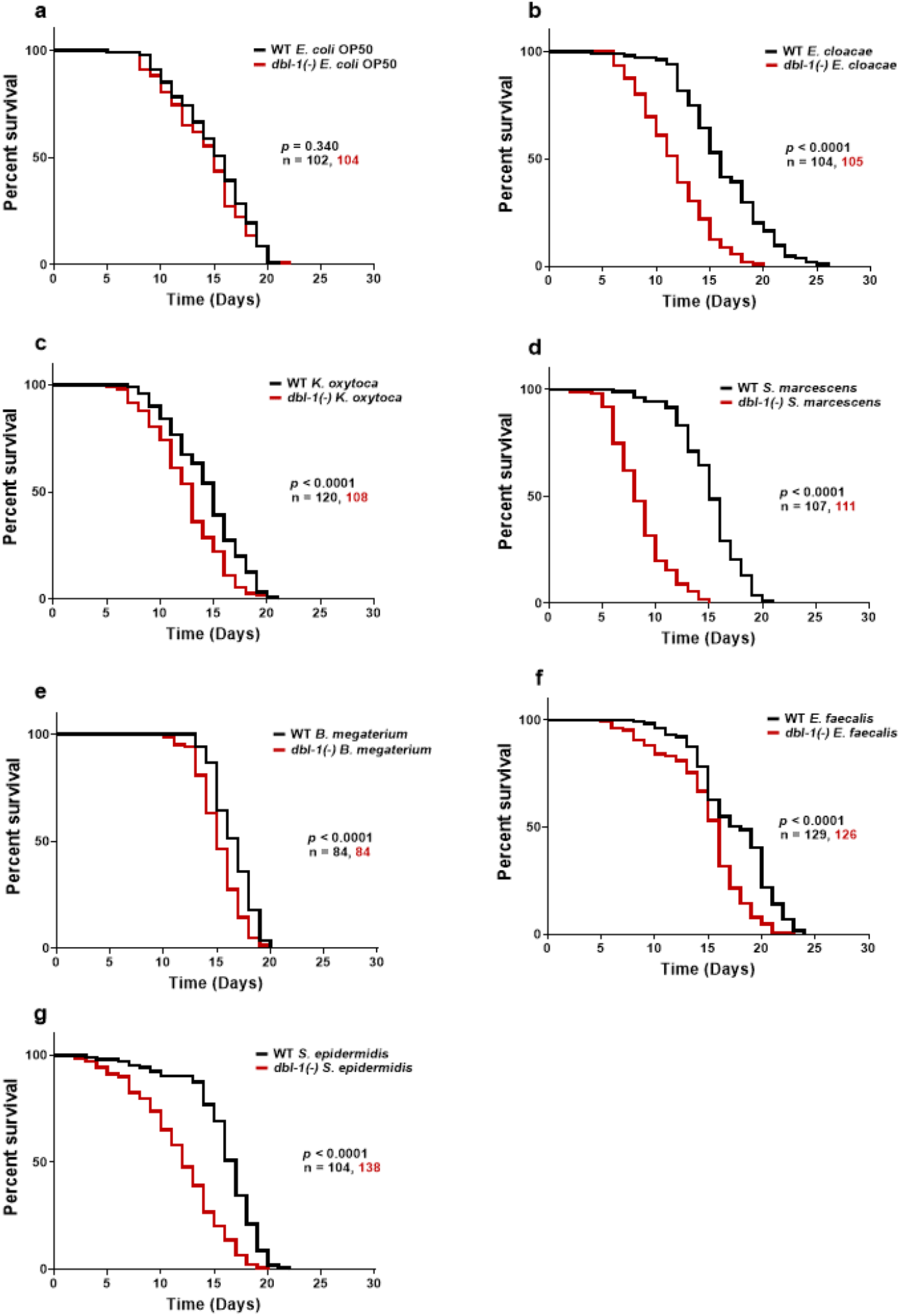
Loss of DBL-1 decreases lifespan of animals exposed to Gram-negative and Gram-positive bacteria. Wild-type and *dbl-1(−)* animals were scored for survival over time from the L4 stage (t=0 hours) on the following bacteria: a) *E. coli* OP50 (control), *E. cloacae*, c) *K. oxytoca*, d) *S. marcescens*, e) *B. megaterium*, f) *E. faecalis*, and g) *S. epidermidis*. Survival fraction was calculated by the Kaplan-Meier method. *p*-values were calculated using log-rank test and *p* <0.01 compared to wild-type animals exposed to the same bacteria was considered significant. One representative trial of at least three is presented.

Loss of DBL-1 has previously been shown to reduce lifespan of animals exposed to fungus *D. coniospora*, Gram-negative strains *E. coli* and *S. marcescens* (Db11), and Gram-positive *E. faecalis*^13,20,41^. To determine if DBL-1 is required in maintaining lifespan of animals subjected to our bacterial panel, we compared lifespans of *dbl-1(−)* animals exposed to the test or control bacteria to the wild type. In our conditions, loss of DBL-1 does not alter lifespan of animals fed on *E. coli* (Fig.1a). However, loss of *dbl-1* results in a significantly shortened lifespan on *S. marcescens*, consistent with a previous report that used the more virulent *S. marcescens* strain Db11 (Fig.1d, Supplementary Table 3)^20^. *dbl-1* mutant animals did not have the lifespan extension seen in wild-type populations on *E. cloacae* or *K. oxytoca*: lifespans of *dbl-1* mutant populations were the same on these two Gram-negative bacterial strains as on *E. coli* (Fig.1b, c, Supplementary Table 3). The lifespan of *dbl-1(−)* animals was extended upon exposure to Gram-positive *B. megaterium* and *E. faecalis* compared to the *E. coli*-fed population’s lifespan, but was not as extended as the wild-type lifespan (Fig.1e, f, Supplementary Table 3). Lastly, *dbl-1* mutant animals displayed a significantly decreased lifespan on *S. epidermidis* (Fig.1g, Supplementary Table 3).

In conclusion, we have identified a panel of human opportunistic pathogens that can be used to interrogate genetic contributions to innate immune defenses. DBL-1 is required for normal lifespan responses regardless of bacterial Gram nature (Fig.1b-g). These results provide evidence that DBL-1 signaling normally confers protection against these bacteria and may play a role in the lifespan extension observed on most bacteria in the panel.

### Loss of DBL-1 and exposure to specific bacteria reduce feeding

One reason for the increased lifespan of *C. elegans* on select bacteria could be that the animals experience dietary restriction because they reduce bacterial consumption^42,43^. To determine if animals reduce feeding on bacteria that increase lifespan, we measured and compared pharyngeal pumping rates of wild-type animals on control and test bacteria. Wild-type animals fed on *E. cloacae* exhibited a small but significant decrease in pharyngeal pumping. Pharyngeal pumping was not significantly reduced after exposure to *K. oxytoca*, which is not consistent with the mild lifespan extension. Wild-type animals fed on *S. marcescens* have the same pumping rate as on the control bacteria (Fig.2a). For wild-type animals fed on the three Gram-positive bacteria, though, the pumping rate was dramatically decreased, consistent with the lifespan extension these strains conferred to wild-type *C. elegans* (Fig.2b).

**Figure 2.**
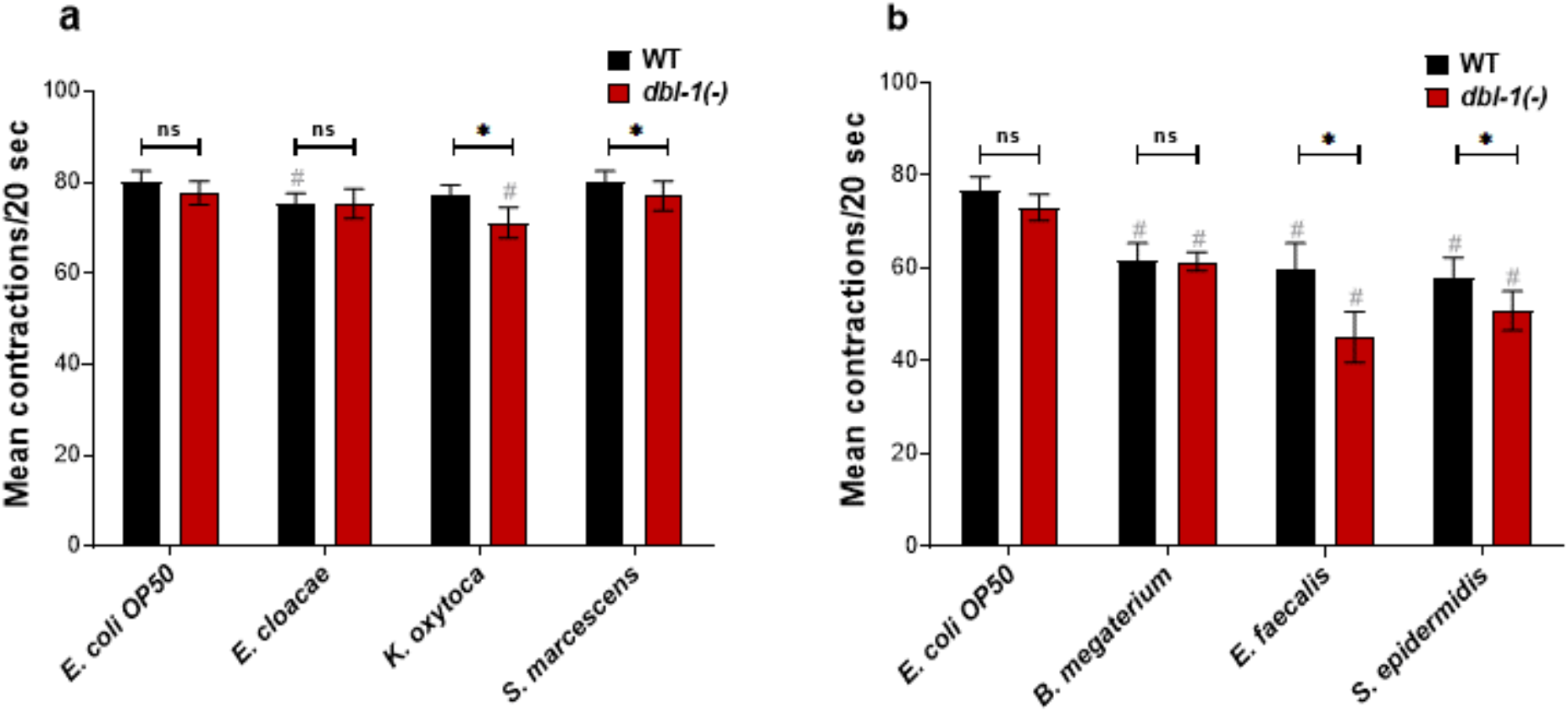
Loss of DBL-1 and exposure to specific bacteria results in decreased pharyngeal pumping. Wild-type and *dbl-1(−)* animals were exposed to the following bacteria: a, b) *E. coli* OP50 (control); a) *E. cloacae*, *K. oxytoca*, *S. marcescens*; b) *B. megaterium*, *E. faecalis*, or *S. epidermidis* at the L4 stage. After 48 hours of exposure, the number of pharyngeal pumps were counted twice per 20 seconds. The pharyngeal pumps were averaged for each animal. One representative trial of at least three is presented. Error bars represent standard deviation. n=12 per condition. * *p* <0.01, ns not significant, compared to wild-type animals exposed to the same bacteria, and # *p* <0.01, respective genotype exposed to test bacteria in comparison to control bacteria by two-way ANOVA followed by unpaired *t*-test.

To determine whether DBL-1 affects feeding response on our panel of test bacteria, we compared the pharyngeal pumping rate of wild-type and *dbl-1(−)* animals on these bacteria. Loss of DBL-1 does not alter the pharyngeal pumping rate of animals fed on the control bacteria (Fig.2a, b). *dbl-1(−)* animals fed on *E. cloacae* do not alter the pumping rate compared to either wild-type animals on *E. cloacae* or *dbl-1* animals on *E. coli* (control). Animals lacking DBL-1 show a mild but significant decrease when they are fed on *K. oxytoca* and *S. marcescens* (Fig.2a). The feeding rate of *dbl-1(−)* animals is further reduced from the wild-type rate on *E. faecalis* and *S. epidermidis* (Fig.2b). There is no reproducibly significant decrease in the pharyngeal pumping rate of *dbl-1(−)* animals fed on *B. megaterium* in comparison to that of the wild type (Fig.2b). These results collectively indicate that while the feeding reduction caused by exposure to these Gram-positive bacteria is independent of DBL-1, a stronger pharyngeal pumping depression in *dbl-1(−)* populations occurs in response to some bacteria (both Gram-negative and -positive bacteria), providing support to the idea that loss of DBL-1 sensitizes animals to certain pathogenic stressors. These findings suggest that even though the organismal responses (lifespan) requiring DBL-1 are similar, the underlying causes might be different, differences in animal feeding being one.

### Intestinal integrity of animals is not altered by loss of *dbl-1* or exposure to specific bacteria

The integrity of intestine can also be disrupted by exposure to pathogenic bacteria, but progressive loss of intestinal integrity is also a feature of aging^33,44^. To further examine the role of DBL-1 in the organismal responses to this panel of bacteria, we investigated the integrity of the intestinal barrier of animals fed on different bacteria. Using a cell-impermeable blue dye, we compared the intestinal barrier function of wild-type and *dbl-1(−)* animals exposed to control and test bacteria when 50% of the population on the test bacteria was still alive. Consistent with previous reports, wild-type animals on *E. coli* exhibited an age-dependent reduction of intestinal integrity^44^. Animals lacking DBL-1 and grown on *E. coli* also displayed a significant decline in intestinal integrity similar to the wild type (Supplementary Fig.1a). Exposure of wild-type or *dbl-1* mutant animals to any of the Gram-negative and -positive bacteria in the panel did not result in further, reproducible decreases of intestinal integrity compared to *E. coli* (Supplementary Fig.1). Therefore, *dbl-1* is not required for intestinal integrity, nor its age-related decline. Furthermore, this result suggests the lifespan changes observed in *dbl-1* mutant populations are not caused by loss of intestinal integrity.

### DBL-1 signaling is required to suppress avoidance against Gram-negative bacteria

*C. elegans* can sense and avoid pathogenic bacteria for protection against harmful environments ^45–47^. To test how our panel of selected Gram-negative and -positive bacteria evokes an avoidance response in *C. elegans*, we performed the avoidance assay. We measured the avoidance response of wild-type animals fed on the test bacteria over the first two days of adulthood and compared it with the avoidance response on the control bacteria. Wild-type animals do not avoid *E. coli* (Fig.3a). We found that wild-type animals mildly avoid *S. marcescens*, but do not avoid *E. cloacae* and *K. oxytoca* (Fig.3b-d). Animals did not avoid the Gram-positive bacteria *E. faecalis* and *S. epidermidis*, but had a moderate to strong response to *B. megaterium*. (Fig.3e-g). This indicates that *S. marcescens*, though it did not extend lifespan, and *B. megaterium* are mildly pathogenic to animals.

**Figure 3.**
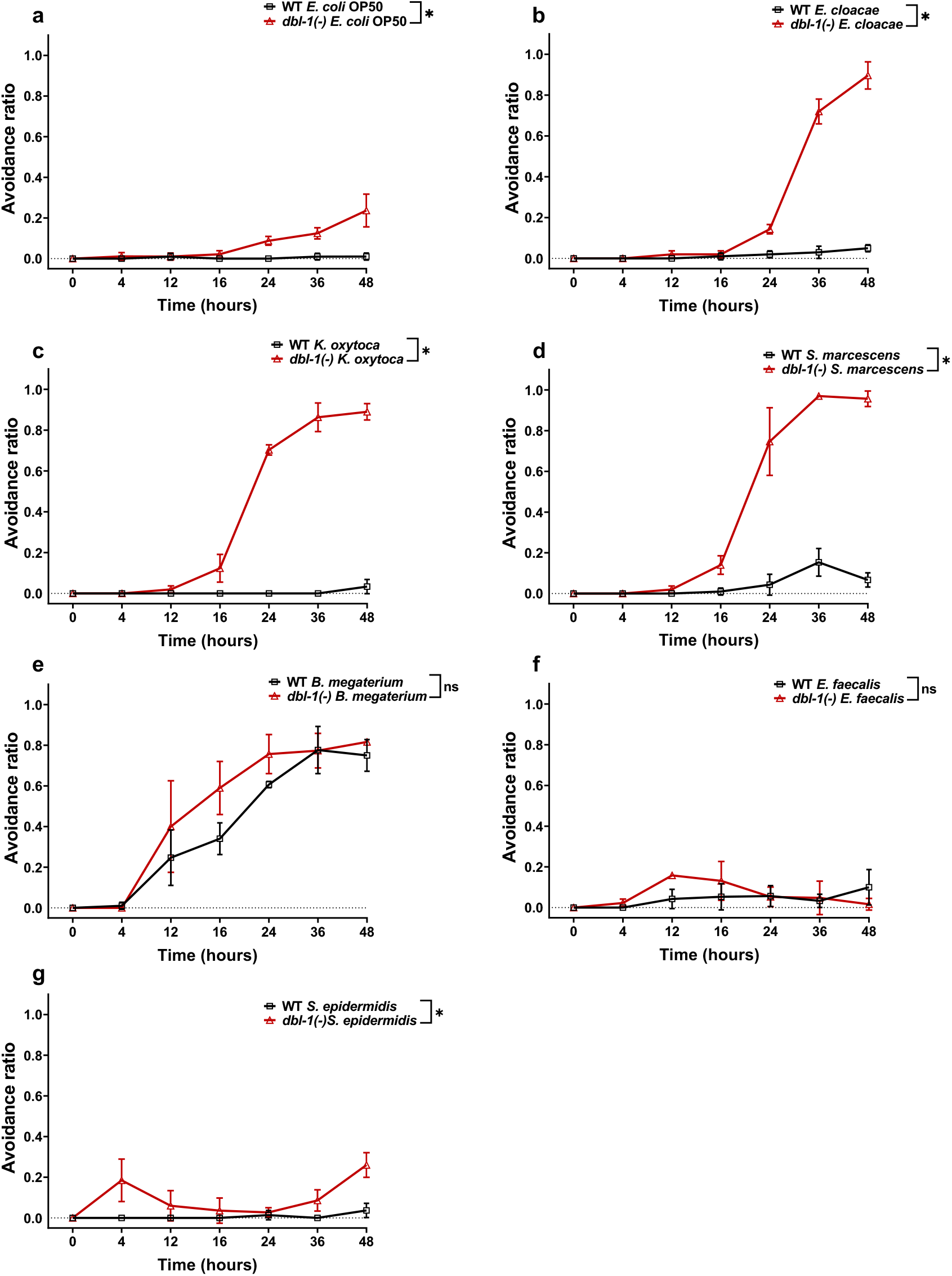
Avoidance to Gram-negative bacteria increases upon loss of DBL-1 signaling. Wild-type and *dbl-1(−)* animals at the L4 stage (t=0 hours) were exposed to the following bacteria: a) *E. coli* OP50 (control), b) *E. cloacae*, c) *K. oxytoca*, d) *S. marcescens*, e) *B. megaterium*, f) *E. faecalis*, or g) *S. epidermidis*. Avoidance of animals to the bacteria was monitored over time. The avoidance ratio was calculated and compared between wild-type and *dbl-1(−)* animals. One representative trial of three is presented. Error bars represent standard deviation. n=30 per condition per trial. * *p* <0.05, and ns not significant, compared to wild-type animals exposed to the same bacteria by repeated measures ANOVA using Tukey’s post-hoc test.

To determine if DBL-1 regulates this avoidance phenotype, we tested avoidance behavior of *dbl-1(−)* exposed to the test bacteria and compared it with *dbl-1(−)* animals fed on the control bacteria. Loss of *dbl-1* usually results in mild but significantly higher avoidance to the control bacteria than the wild type (Fig.3a), in support of previous findings^48^. *dbl-1(−)* animals displayed a striking, strong avoidance response to all three tested Gram-negative bacteria and a mild response to *S. epidermidis* (Fig.3b-d, g). Interestingly, *dbl-1(−)* animals exhibited a similar avoidance behavior in response to *B. megaterium* and *E. faecalis* exposure (Fig.3e, f). This indicates that upon loss of DBL-1, animals perceive *E. cloacae*, *K. oxytoca*, *S. marcescens*, and *S. epidermidis* as more pathogenic. Avoidance responses to *B. megaterium* and *E. faecalis* is independent of DBL-1 levels. These results indicate that DBL-1 normally suppresses avoidance and loss of DBL-1 results in robust avoidance responses that depend on the type of bacterial exposure.

### SMA-4 acts independently of other DBL-1 core signaling components to suppress avoidance to Gram-positive bacteria

Because we observed strong bacteria-specific avoidance responses that were DBL-1-dependent, we next asked if the canonical DBL-1 signaling pathway is required to attenuate this response. Canonical signaling occurs by DBL-1 ligand binding to receptors SMA-6 and DAF-4, which activate downstream transcription factors SMA-2, SMA-3, and SMA-4 ^49^. A non-canonical DBL-1 pathway, which does not signal through SMA-2 and SMA-4, is required for *C. elegans* to respond to the fungus *D. coniospora*^13^. We measured the avoidance response of *sma-2(−)*, *sma-3(−)*, and *sma-4(−)* animals fed on the test Gram-negative or -positive bacteria and compared them with the avoidance response on the control bacteria. On the control bacteria, loss of *sma-3* did not result in significantly increased avoidance. However, populations lacking *sma-2* avoided the control bacteria mildly to moderately, and populations lacking *sma-4* showed a moderate to strong avoidance response (Fig.4a). *sma-2(−)*, *sma-3(−)*, and *sma-4(−)* populations displayed a strong, reproducible avoidance response to all Gram-negative bacterial strains tested that was comparable to the *dbl-1(−)* response (Fig.4b-d). In comparison, the response of Smad mutant populations to the panel of Gram-positive bacteria was notably different. Similar to loss of *dbl-1*, loss of *sma-2* or *sma-3* did not increase avoidance responses to the three Gram-positive bacterial strains. However, loss of *sma-4* resulted in moderate to strong avoidance to all three Gram-positive strains (Fig.4e-g). Our findings indicate that while DBL-1 and SMA-3 do not play a major role in responding to *E. coli*, SMA-2 and SMA-4 are required to suppresses avoidance to this standard lab food. Furthermore, our results support that canonical DBL-1 signaling plays a major role in suppressing animal avoidance responses to Gram-negative bacteria, but is not required for responding to Gram-positive bacteria. Interestingly, because of the strong effect loss of SMA-4 has on avoidance behavior, SMA-4 may act with another factor independent of the DBL-1 signaling pathway that heavily influences avoidance responses to Gram-positive bacteria.

**Figure 4.**
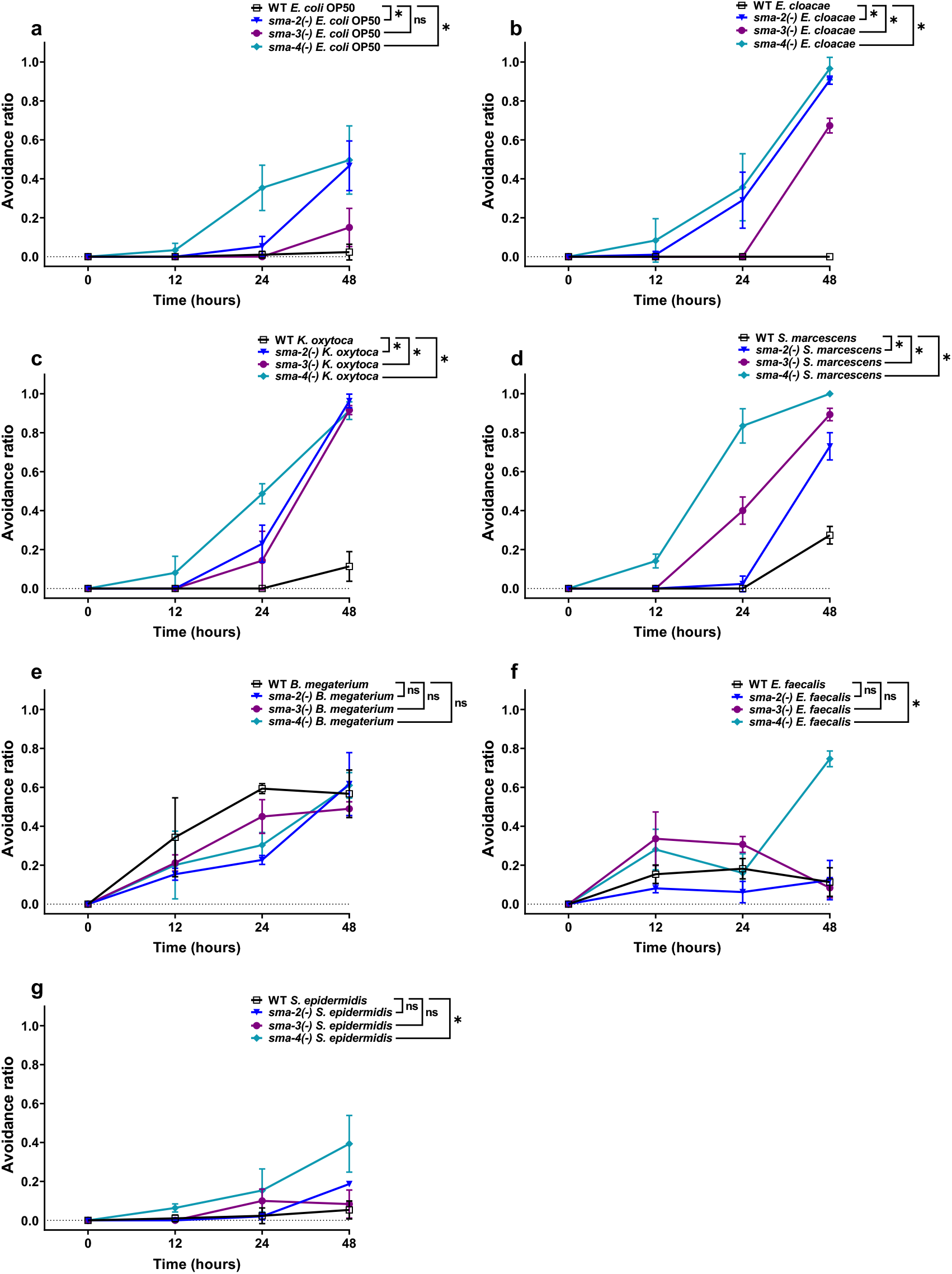
Avoidance to Gram-negative bacteria increases upon loss of canonical DBL-1 signaling. Wild-type, *sma-2(−)*, *sma-3(−)*, and *sma-4(−)* animals at the L4 stage (t=0 hours) were exposed to the following bacteria: a) *E. coli* OP50 (control), b) *E. cloacae*, b) *K. oxytoca*, d) *S. marcescens*, e) *B. megaterium*, f) *E. faecalis*, or g) *S. epidermidis*. Avoidance of animals to the bacteria was monitored over time. The avoidance ratio was calculated and compared between wild-type and Smad mutant animals. Avoidance ratio = number of animals off the bacterial lawn/ total number of animals. One representative trial of three is presented. Error bars represent standard deviation. n=30 per condition per trial. * *p* <0.05, ns not significant, compared to wild-type animals exposed to the same bacteria by repeated measures ANOVA using Tukey’s post-hoc test.

### *sma-4* expression is specifically induced in response to Gram-positive bacteria

We next asked if the different avoidance responses to Gram-negative and -positive bacteria are associated with altered gene expression of the DBL-1 Smads. We tested gene expression levels of *sma-2*, *sma-3*, and *sma-4* in wild-type and *dbl-1* mutant backgrounds in response to our panel of bacteria. With the exception of *S. epidermidis* exposure, the relative levels of *sma-2* mRNA were consistently decreased in the *dbl-1* mutant background, but the test bacterial strains had no effect on *sma-2* expression (Fig.5a-d). The relative levels of *sma-3* were not reproducibly different between *dbl-1* and wild-type backgrounds, and the overall expression of *sma-3* in animals exposed to the test bacterial strains was not altered. *K. oxytoca* and *E. faecalis* conditions did result in significantly increased *sma-*3 levels that were not observed in the *dbl-1* background (Fig.5e-h). On control and Gram-negative bacteria, *sma-4* expression was similar in the *dbl-1* background as in the wild type. In addition, *sma-4* expression was not changed by exposure to the test Gram-negative bacteria (Fig.5i, k). However, *sma-4* was significantly induced in response to all three Gram-positive bacteria in both wild-type and *dbl-1* mutant backgrounds, albeit less in the *dbl-1(−)* populations (Fig.5j, l). Together, these results suggest that the DBL-1 Smads are differently regulated at the level of gene expression by molecular pathways that are responsive to specific bacterial challenges. *sma-2*, but not *sma-3*, requires DBL-1 for full expression regardless of bacterial food source. Neither *sma-2* nor *sma-3* expression is largely affected by test bacteria. In contrast, *sma-4* expression is responsive to Gram-positive bacteria but not Gram-negative bacteria in the panel, possibly by both DBL-1 and DBL-1-independent mechanisms.

**Figure 5.**
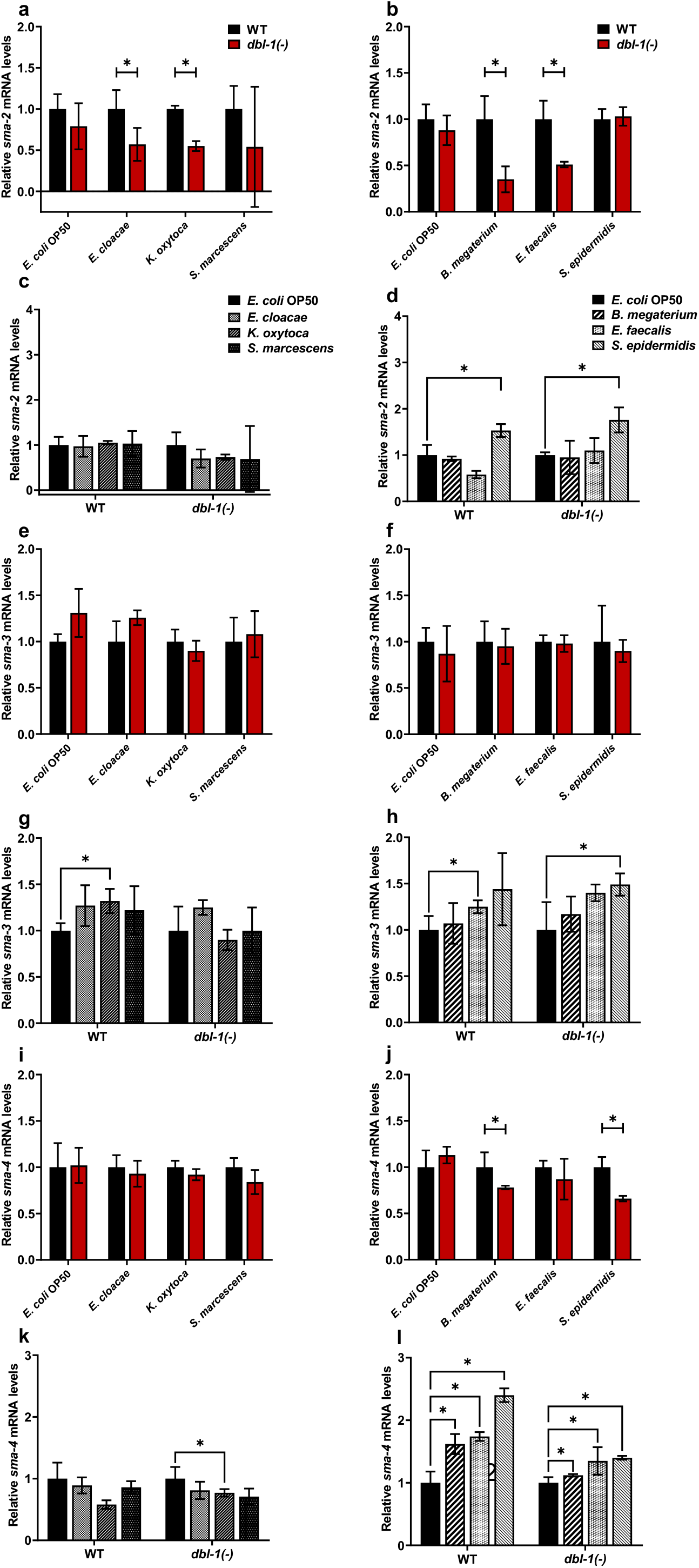
Smad transcription factors gene expression is altered by specific bacteria. Wild-type and *dbl-1(−)* animals at the L4 stage were exposed to *E. coli* OP50 (control), *E. cloacae*, *K. oxytoca*, *S. marcescens*, *B. megaterium*, *E. faecalis*, or *S. epidermidis* for 48 hours. mRNA expression levels of a-d) *sma-2*, e-h) *sma-3*, and i-l) *sma-4* were quantitated by real-time PCR. Experiments were performed in triplicate and in three independent trials. One representative trial of three is presented. Error bars represent standard deviation. * *p* <0.05 in a, b, e, f, i, j, mRNA expression level in *dbl-1(−)* population compared to wild-type population exposed to the same bacteria by unpaired *t*-test. * *p* <0.05 in c, d, g, h, k, l, mRNA expression level in respective genotype exposed to test bacteria compared to control bacteria by unpaired *t*-test.

### DBL-1 signaling is activated in response to Gram-negative bacteria and is repressed in response to Gram-positive bacteria

Because we observed changes in expression levels of the Smads, we asked if DBL-1 signaling activity is altered in response to the bacteria panel. To address this question, we challenged wild-type or *dbl-1(−)* animals expressing an integrated fluorescent DBL-1 pathway reporter to these bacteria and analyzing reporter fluorescence in L2 hypodermal nuclei. The expression of this reporter is robust in hypodermal nuclei at the L2 stage, and changes in DBL-1 affect hypodermal expression of RAD-SMAD^39^. This reporter (called RAD-SMAD) consists of the GFP gene under the control of multiple copies of a Smad binding element sequence^50^. In *dbl-1(−)* animals on *E. coli*, RAD-SMAD fluorescence was not detectable or very faint. In response to Gram-negative bacteria, reporter fluorescence in the wild-type background was significantly (two- to four-fold) increased compared to control bacterial conditions (Fig.6a-d, h). Reporter intensity in *dbl-1(−)* animals on all Gram-negative bacteria remained either undetectable or very faint. In stark contrast, RAD-SMAD hypodermal fluorescence in wild-type animals was lost upon exposure to Gram-positive *S. epidermidis* (94% (n=66) had no detectable expression) and *B. megaterium* (100% (n=42)). In about half of the wild-type population fed *E. faecalis*, hypodermal fluorescence was not detected (46% (n=53)), but fluorescence levels were wild type in those animals expressing RAD-SMAD in the hypoderm (Fig.6e-g, Supplementary Table 4). For animals lacking *dbl-*1 fed on any of the Gram-positive bacterial conditions, hypodermal RAD-SMAD fluorescence was undetectable or very faint. Exposure of *dbl-1(−)* animals to *S. epidermidis* and *B. megaterium* resulted in a stronger repression of RAD-SMAD activity than exposure to *E. faecalis*, similar to the wild-type responses to these Gram-positive bacteria (Supplementary Table 4). In general, these results indicate that animals induce a DBL-1 signaling response to Gram-negative bacteria but repress DBL-1 signaling in response to Gram-positive bacteria. Furthermore, DBL-1 signaling levels appear to be modulated depending on the specific bacterial challenge that animals encounter.

**Figure 6.**
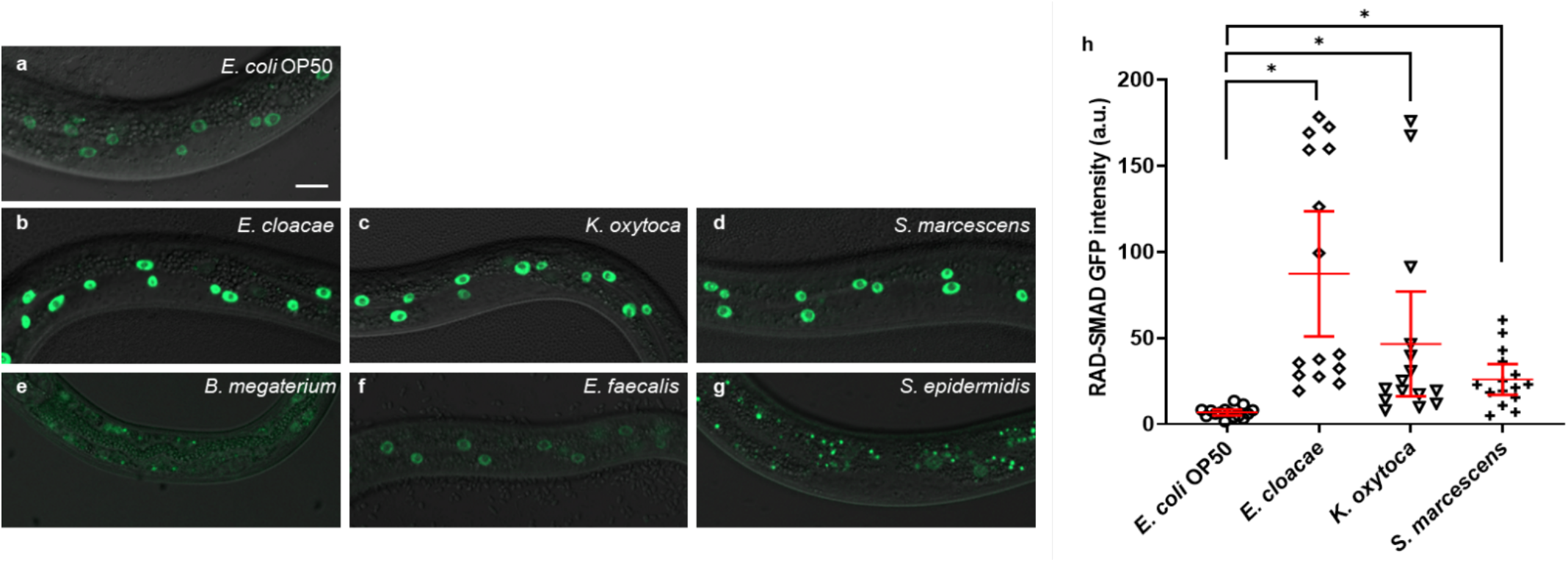
DBL-1 signaling is activated upon exposure to Gram-negative bacteria but is repressed in response to Gram-positive bacteria. L4 animals expressing the RAD-SMAD reporter in a wild-type background were exposed to a, h) *E. coli* OP50 (control), b, h) *E. cloacae*, c, h) *K. oxytoca*, d, h) *S. marcescens*, e) *B. megaterium*, f) *E. faecalis*, or g) *S. epidermidis* and L2-stage progeny were imaged. Mean RAD-SMAD fluorescence intensity of five hypodermal nuclei per animal was quantitated and compared. Experiments were performed in three independent trials. One representative trial is presented. Error bars represent 95% confidence intervals. n=15 per condition in each trial. * *p* <0.05, mean fluorescence intensity in wild-type background on test bacteria compared to control bacteria by unpaired *t*-test. Scale bar, 10 μm.

### DBL-1 mediates both common and specific gene expression responses to Gram-negative and -positive bacteria

We then asked if this differential modulation of DBL-1 signaling activity translated to bacterial-specific downstream transcriptional responses. To identify the role of DBL-1 in differentially regulating transcription of downstream genes, we performed RNA sequencing using wild-type and *dbl-1(−)* animals exposed to the control or test bacteria (Gram-negative *S. marcescens* or Gram-positive *E. faecalis*). The animals were synchronized as L4s and fed on the control or test bacteria for 48 hours before analysis. In animals lacking DBL-1 and fed control bacteria, 83 genes were down-regulated and 49 genes were upregulated compared to the wild type (*p* < 0.01, Supplementary Fig.2a). Some genes and gene classes previously reported to be regulated by DBL-1 at different developmental stages were also regulated by DBL-1 in two-day adults^20,24,25,51^. In *dbl-1(−)* animals fed on *S. marcescens*, 102 genes were down-regulated and 117 genes were upregulated (Supplementary Fig.2b). In *dbl-1(−)* animals fed on *E. faecalis*, 63 genes were down-regulated and 64 genes were upregulated compared to the wild type (Supplementary Fig.2c). The lower number of highly regulated genes between wild-type and *dbl-1(−)* animals fed on *E. faecalis* is consistent with the reduced DBL-1 reporter fluorescence—and therefore DBL-1 pathway signaling—in wild-type animals on *E. faecalis* (Fig.6, Supplementary Fig.2). Notably, some highly regulated genes were common in response to both pathogenic bacterial strains, but some genes that were differentially regulated by DBL-1 were unique in response to either *S. marcescens* or *E. faecalis* exposure. Using WormCat, gene enrichment analysis of genes regulated by DBL-1 in response to *S. marcescens* or *E. faecalis* exposure revealed differential regulation of sets of genes involved in pathogen response, stress response, lipid metabolism, and transmembrane transport (Supplementary Data 1)^52^.

We focused on the DBL-1-regulated genes that have known or putative roles in innate immunity. These genes were induced in wild-type animals upon exposure to *S. marcescens* or *E. faecalis* and this induction was lost in *dbl-1(−)*. We also found some genes to be upregulated upon loss of *dbl-1* in response to *S. marcescens* or *E. faecalis*. Some gene classes are highly regulated in response to these bacteria, but the specific genes within these families differed, including lysozyme, aspartyl protease, saposin-like, and C-type lectin genes (Fig.7). These results suggest that DBL-1 is involved in regulating (positively and negatively) transcription of some innate immunity genes specific to the bacterial exposure and some genes that are commonly regulated upon exposure to different bacteria. This supports a role for DBL-1 signaling in differentially regulating host responses to bacteria.

**Figure 7.**
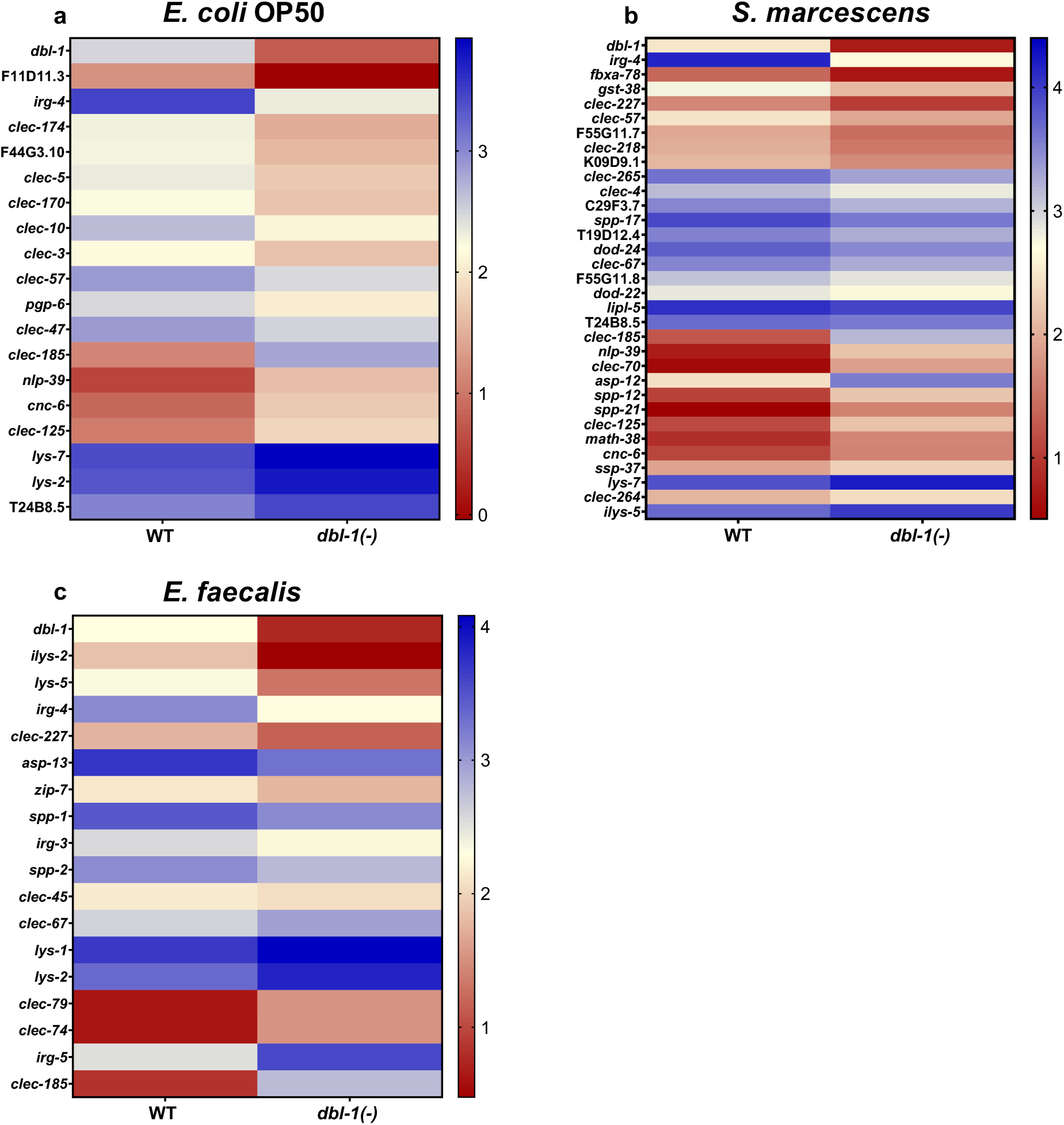
Expression of innate immunity genes is differentially regulated by DBL-1 signaling in different bacterial environments. L4 wild-type and *dbl-1(−)* animals were exposed for two days to *E. coli* OP50 (control), *S. marcescens*, or *E. faecalis*. RNA-seq analysis reveal significant changes (adjusted *p*-value <0.01) in gene expression of animals lacking DBL-1. Heatmaps show differential innate immunity gene expression in animals lacking DBL-1 exposed to a) *E. coli* OP50, b) *S. marcescens*, and c) *E. faecalis* in comparison to wild-type animals exposed to the same bacteria. Average log FPKM values from three independent trials are represented.

### DBL-1 differentially regulates expression of innate immunity genes specific to the Gram nature of bacteria

From the RNA-sequencing results, we identified a panel of DBL-1 responsive innate immune genes that displayed differential responses to pathogen exposure. To determine if the expression of candidate target immunity genes is regulated by DBL-1 signaling in response to a wider variety of bacterial exposures, we used reporters of select immunity-related genes. Based on the RNA sequencing results, we selected *dod-22*, F55G11.7, *irg-4*, and *dod-24*. We also selected *ilys-3*, a known Gram-positive-responsive gene^9^. We tested expression of transcriptional reporters of these genes upon exposure to different Gram-negative and -positive bacteria. We measured and compared expression of these reporter genes in wild-type and *dbl-1(−)* backgrounds exposed to control or test bacteria. Animals were synchronized as L4s and fed on control, Gram-negative, or Gram-positive bacteria for 48 hours. Basal expression of these selected genes was measured in animals fed on the control *E. coli*.

*dod-22* is a gene that is known to be regulated by the insulin-like signaling pathway transcription factor, DAF-16 ^21^. It is known to be involved in defense response to Gram-negative bacteria^16^. The *dod-22* reporter is induced in the wild-type background in the presence of all Gram-negative test bacteria compared to the control, but is not induced in response to the panel of Gram-positive bacteria. This induction of the *dod-22* reporter on Gram-negative bacteria is partly lost in the *dbl-1(−)* background, though loss of *dbl-1* does not affect expression levels on the control *E. coli*. *B. megaterium* does not lead to a reproducible reduction of *dod-22* reporter fluorescence in the wild-type background, but a further reduction of fluorescence is observed in the *dbl-1(−)* background (Fig.8a, b). These results indicate that DBL-1 is not required for the basal expression of *dod-22*, but is required for *dod-22* induction on Gram-negative bacteria.

**Figure 8.**
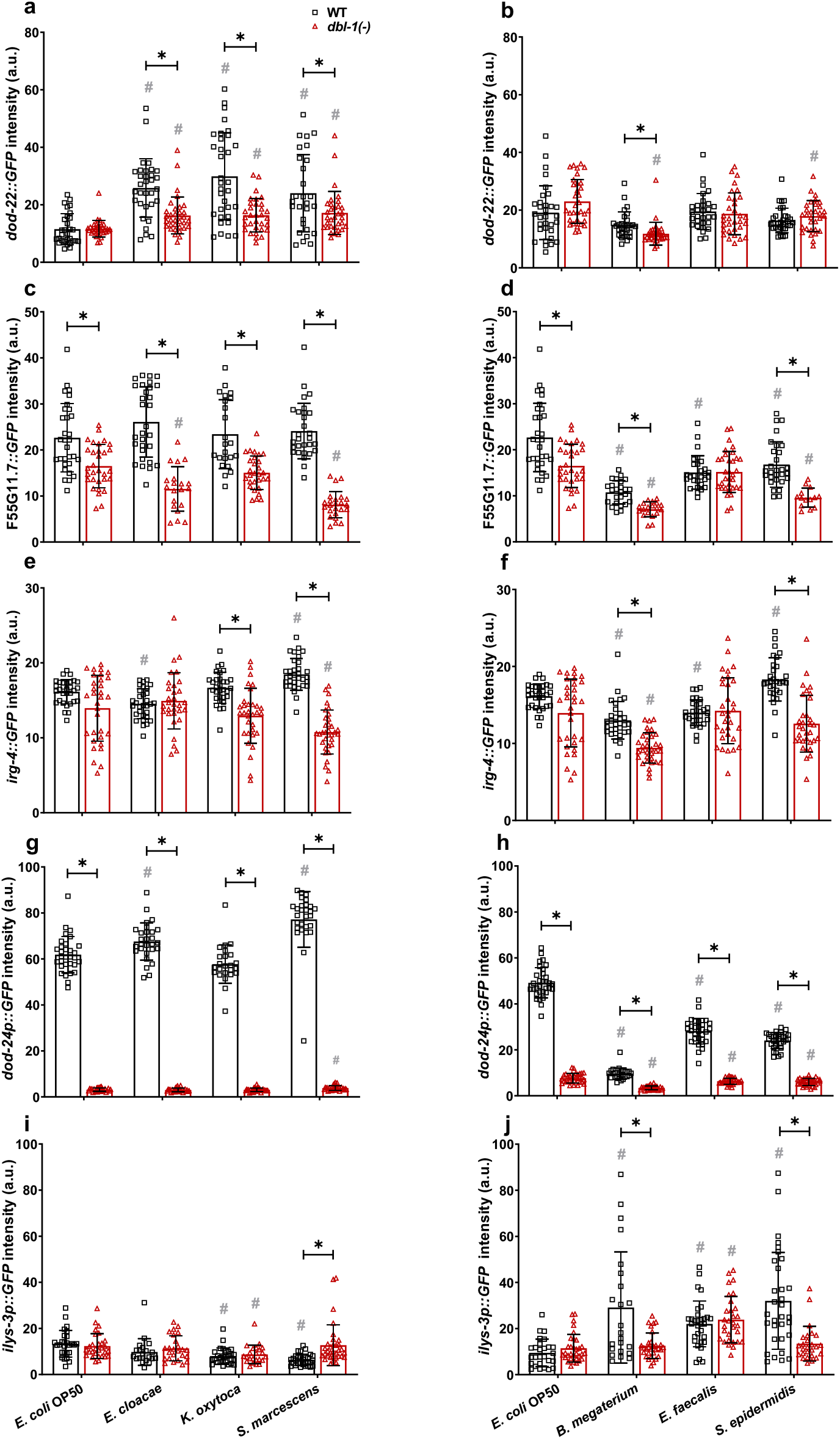
Innate immune reporter activity is regulated by exposure to specific bacteria and by DBL-1 signaling. Comparison of a, b) *dod-22::*GFP, c, d) F55G11.7::GFP, e, f) *irg-4::*GFP, g, h) *dod-24p::*GFP, and i, j) *ilys-3p::*GFP intensities in adult wild-type and *dbl-1(−)* animals after a two-day exposure to the following bacteria; control *E. coli* OP50, *E. cloacae*, *K.* oxytoca, *S. marcescens, B. megaterium*, *E. faecalis*, or *S. epidermidis*. Imaging conditions including exposure times were consistent with respective control. Three independent trials were performed. One representative trial is shown. Error bars represent standard deviation. n= at least 15 per condition in each trial. * *p* <0.05 compared to wild-type animals exposed to the same bacteria and # *p* <0.05 respective genotype exposed to test bacteria in comparison to control bacteria, by two-way ANOVA followed by unpaired *t*-test.

F55G11.7 is involved in innate immune responses to both Gram-negative and -positive bacteria in *C. elegans* and has been shown to be regulated by DAF-16/insulin, MAPK, and DBL-1 signaling pathways^16^. Expression of the F55G11.7 reporter did not change upon exposure to all test bacteria in the wild-type background except *B. megaterium*, where reporter expression was reduced (in two of three trials). We observed a significant reduction of F55G11.7 reporter activity in animals lacking DBL-1 except in response to *E. faecalis*. Additionally, a further reduction of F55G11.7 reporter fluorescence in *dbl-1(−)* animals was observed in response to Gram-positive *B. megaterium* and *S. epidermidis* compared to the response on control bacteria (Fig.8c, d). These findings indicate that expression of F55G11.7 is not altered upon exposure to most of the test bacteria but is generally down-regulated upon loss of DBL-1, in contrast with previous findings^16^.

*irg-4* is known to be involved in defense response to Gram-negative bacteria and has been shown to be regulated by DAF-16/insulin, MAPK, and DBL-1 signaling pathways^53–56^. In the wild-type background, we observed a visible, reproducible induction of *irg-4* reporter activity in response to Gram-negative *S. marcescens*, but not to *K. oxytoca* or *E. cloacae*. Expression of the *irg-4* reporter in wild-type animals mildly increased in response to *S. epidermidis*, but not the other Gram-positive bacteria reproducibly. Loss of DBL-1 did not reproducibly alter *irg-4* reporter activity in control conditions. However, *irg-4* reporter induction in response to Gram-negative *S. marcescens* and Gram-positive *S. epidermidis* was lost in the *dbl-1(−)* background. (Fig.8e, f). Indeed, *irg-4* reporter expression was reduced compared to the wild-type background on all bacteria except *E. cloacae* and *E. faecalis*. These results indicate that *irg-4* is responsive to a broad range of bacteria and is regulated in part by DBL-1 signaling.

*dod-24*, which is regulated by the insulin-like signaling transcription factor DAF-16, is involved in defense response to Gram-negative bacteria^53^. We observed robust expression of *dod-24* reporter activity in all tested Gram-negative bacteria, including the control, *E. cloacae*, *K. oxytoca*, and *S. marcescens*. Additionally, we observed further induction in response to *E. cloacae* and *S. marcescens*. We observed a striking decrease of *dod-24* reporter activity in wild-type animals exposed to all tested Gram-positive bacteria including *B. megaterium*, *E. faecalis*, and *S. epidermidis*. Loss of DBL-1 resulted in a significant reduction of *dod-24* reporter activity in control conditions. *dod-24* reporter activity was also drastically reduced in all Gram-negative bacterial conditions to levels significantly lower than the wild type (Fig.8g). In the three Gram-positive conditions, loss of DBL-1 resulted in a further decrease of *dod-24* reporter fluorescence relative to the wild type (Fig.8h). These results confirm that *dod-24* is differentially expressed in response to an array of Gram-negative and -positive bacteria and indicate that DBL-1 signaling plays a major role in regulating *dod-24* expression.

*ilys-3* exhibits lysozyme activity and is involved in responding to Gram-positive bacteria^9^. *ilys-3* reporter activity in wild-type animals was unchanged or reduced in response to the tested Gram-negative bacteria (Fig.8i). We observed induction of *ilys-3* reporter activity upon exposure to all tested Gram-positive bacteria including *B. megaterium*, *E. faecalis*, and *S. epidermidis* (Fig.8j). Loss of DBL-1 did not alter *ilys-3* reporter activity in control conditions. The *ilys-3* reporter activity remained at relatively low levels in animals lacking DBL-1 exposed to Gram-negative bacteria (Fig.8i). However, *ilys-3* reporter activity also remained at relatively low levels upon loss of DBL-1 in response to Gram-positive bacteria *B. megaterium* and *S. epidermidis*, but was wild type in response to *E. faecalis* (Fig.8j). These results suggest that while DBL-1 is not required for basal levels of *ilys-3* expression, it is required for the induction of *ilys-3* expression in response to some Gram-positive bacteria.

These results collectively indicate the specific role of DBL-1 signaling in antimicrobial gene expression that helps tailor defense responses to specific bacterial challenges.

## DISCUSSION

Animals are subjected to a range of bacterial challenges, and how they respond is critical to the animals’ health. Understanding how hosts respond to different pathogens is important for developing therapeutic strategies to help fight infections and prevent diseases. Dissecting the cell signaling pathways involved in host responses and their roles is critical. Our work expands the current understanding of how an organism integrates an arsenal of responses—from the molecular to the organismal—to different bacteria, and identifies role of the DBL-1/TGF-β pathway in robust host-specific responses to different types of bacteria. For this work, we established a panel of human opportunistic pathogens, including three Gram-negative and three Gram-positive strains, for the study of long-term innate immune responses in the roundworm *C. elegans*. The bacteria we selected for the panel elicit unique host response patterns that allowed us to interrogate the role of the DBL-1 pathway in responding to different bacterial exposures. While DBL-1 has a known role in transcriptionally regulating innate immune gene expression, we show that the specific responses mediated by DBL-1 are not only molecular, but are also behavioral.

Our results support a model that the DBL-1 signaling pathway influences the organisms’ perception of pathogenicity and helps keep the innate immune responses in check. Animals with reduced DBL-1 signaling—whether by downregulating signaling or by mutation—perceive the environment as more threatening and respond accordingly. DBL-1 pathway mutants display an outsized avoidance response to the Gram-negative bacteria. These animals also reduce intake of select Gram-negative bacteria as yet another way to reduce animals’ interaction with the threat. Animals lacking DBL-1 also display a reduced lifespan in response to the Gram-negative bacteria that are sensed to be more pathogenic. There is also a correlation between the reduced feeding and extension of lifespan observed in animals exposed to select bacteria. Because exposure to the Gram-positive bacteria reduce the DBL-1 signaling activity, the avoidance response did not alter dramatically upon loss of DBL-1. The intake of the Gram-positive bacteria was reduced in both wild-type and *dbl-1(−)* populations which correlated with extended lifespan observed for both populations on the Gram-positive bacteria. Damage to the intestine was not the underlying cause for the DBL-1-mediated lifespan alterations in response to the tested bacteria. Determining how DBL-1 is involved in such behavioral modifications in response to different bacteria warrants future investigation.

DBL-1 signaling is also important for molecular responses to both Gram negative and Gram-positive bacteria. Regulation of DBL-1 signaling is part of the host’s molecular response: the Gram-negative bacteria of our panel induced DBL-1 signaling while the Gram-positive bacteria repressed DBL-1 signaling. However, within these two bacterial groups, the host responses were tailored to the specific bacterial challenge. Our results show that DBL-1 signaling is also an important part of this molecular antimicrobial “fingerprint”^16^. Our RNA-sequencing analyses indicate both common and unique transcriptome-wide alterations mediated by DBL-1 after two days of exposure to Gram-negative and Gram-positive bacteria. We find that DBL-1 signaling is involved in activating as well as repressing innate immunity genes to maintain a balance of host immune responses (to possibly avoid overactivation of host immunity). In response to our panel of Gram-negative bacteria, DBL-1 signaling activity was induced and it further regulated expression of unique downstream innate immunity genes. In contrast, even though the DBL-1 signaling activity was repressed in response to the tested Gram-positive bacteria, DBL-1 was required to regulate expression of target immunity genes. While many gene classes differentially regulated by specific pathogens have been previously identified as important innate immune response genes, our work highlights the role that DBL-1 plays in tailoring the molecular responses *C. elegans* engages against a range of pathogens^9,11,16,53,54,57^.

Another major finding of this work is the differential requirement of the SMAD machinery to mediate avoidance responses to our panel of bacteria. Olofsson previously showed that loss of DBL-1, SMA-2, or SMA-4 increases avoidance of *E. coli*^48^. While our results with *sma-2* and *sma-4* are similar to theirs, we observe only mild avoidance of *E. coli* upon loss of DBL-1^48^. However, our work demonstrates that canonical DBL-1 signaling strongly suppresses avoidance to Gram-negative bacteria, but not to Gram-positive bacteria (Fig.3). *sma-4* mutants generally displayed stronger avoidance responses to both control and test bacteria suggesting a DBL-1-independent role for SMA-4 in suppressing avoidance responses. SMA-4 appears to play a double role in innate immune responses. Its starring role in immunity is with the DBL-1 pathway. DBL-1-independent induction of *sma-4* in response to Gram-positive bacterial conditions was also observed. This specificity of *sma-4* induction by the Gram-positive bacteria indicates specificity of innate immune responses. These results indicate that SMA-4 is not only recruited for defenses by something other than DBL-1, but also acts independently of the core DBL-1 pathway. PMK-1/MAPK signaling may be involved in regulating it as SMA-4 is predicted to genetically interact with PMK-1 ^58^. Interestingly, ATF-7, a transcription factor activated by PMK-1, is required for downregulation of *sma-4*—but not other DBL-1 pathway component genes—in wild-type animals exposed to Gram-negative *P. aeruginosa* PA14 ^59^. It will be of interest to discover if SMA-4 plays a broader role in the innate immune response than acting in the DBL-1 pathway.

Overall, we propose that loss of DBL-1 signaling changes the animal’s perception of the environment as more hostile, and this results in more robust protective responses that depend on the specific bacterial challenge. This may help explain the neuronal source of DBL-1 secretion that then targets hypoderm, intestine, and pharynx. DBL-1 secreted from the AVA interneurons activates DBL-1 signaling in the hypodermis to regulate aversive learning upon exposure to Gram-negative *P. aeruginosa*, but the neuronal circuit(s) used by DBL-1 to direct aversive behaviors remains to be identified^27^. SMA-4 plays a double role in innate immune responses, acting as part of the core DBL-1 signaling pathway but also acting in another additive way, suggesting crosstalk with other signaling pathways. Several studies report roles of signaling pathways including MAPK and insulin-like signaling pathways in mounting immune responses to many Gram-positive bacteria. Future work may identify the possible crosstalk mechanisms with the DBL-1 pathway in regulating organismal defense responses. In summary, these findings support a central role for DBL-1/TGF-β signaling not only in crafting tailored responses to unique bacterial challenges, from transcription of specific innate immunity genes to behavioral responses but also being modulated in response to bacteria.

## ACKNOWLEDGEMENTS

We thank M. Farhan Lakdawala and Rosylin Roy for technical assistance. We thank James Lundgren, Paul Yeatts, and TWU’s Center for Research Design and Analysis for assistance with statistical analyses. Some bacterial strains were provided by Amy Jo Hammett. Some strains were obtained from the *Caenorhabditis* Genetics Center (CGC), which is funded by NIH Office of Research Infrastructure Programs (P40 OD010440). We thank WormBase. We thank Laura Hanson and Gumienny lab members for constructive feedback. Novogene provided Supplementary Figure 2. This work was supported by NIH R01GM097591 to TLG, TWU Research Enhancement Program funding to T.L.G., internal funding by Texas Woman’s University to T.L.G., and TWU Experiential Learning Scholar Awards to B.M.

## AUTHOR CONTRIBUTIONS

B.M. and T.L.G. conceptualized and designed experiments, interpreted data, and contributed to writing; B.M. performed experiments.

