## Supplemental tables and figures for "The DBL-1/TGF-β signaling pathway regulates pathogen-specific innate immune responses in *C. elegans*"

Supplementary Table 1. List of strains

Strains used include:

| Strain | Genotype |
|---|---|
| N2 | Wild type |
| NU3 | *dbl-1(nk3)* V (referred to as *dbl-1(-)* in this work) |
| CB502 | *sma-2(e502)* III (referred to as *sma-2(-)* in this work) |
| CB491 | *sma-3(e502)* III (referred to as *sma-3(-)* in this work) |
| DR1369 | *sma-4(e729)* III (referred to as *sma-4(-)* in this work) |
| LW2436 | *jjIs2277[pCXT51(5*RLR::pes-10p(deleted)::GFP) + LiuFD61(mec-7p::RFP)]* I or IV (RAD-SMAD) |
| CB6710 | *eEx650[ilys-3p::GFP + unc-119(+)]* |
| CF3556 | *agIs6[dod-24p::GFP]* |
| SAL139 | *denEx17[dod-22::GFP + unc-119(+)]* |
| SAL143 | *denEx21[*F55G11.7*::GFP + unc-119(+)]* |
| SAL148 | *denEx26[irg-4::GFP + unc-119(+)]* |

Strains created for this work include:

| Strain | Genotype |
|---|---|
| TLG803 | *dbl-1(nk3)* V; *agIs6[dod-24p::GFP]* |
| TLG804 | *dbl-1(nk3)* V; *eEx650[ilys-3p::GFP + unc-119(+)]* |
| TLG805 | *dbl-1(nk3)* V; *denEx26[irg-4::GFP + unc-119(+)]* |
| TLG806 | *dbl-1(nk3)* V; *denEx21[*F55G11.7*::GFP + unc-119(+)]* |
| TLG807 | *dbl-1(nk3)* V; *denEx17[dod-22::GFP + unc-119(+)]* |
| TLG810 | *jjIs2277[pCXT51(5*RLR::pes-10p(deleted)::GFP) + LiuFD61(mec-7p::RFP)]* I or IV; *dbl-1(nk3)* V |

Supplementary Table 2. List of primers for qRT-PCR

| Target gene | Forward primer (5'->3') | Reverse primer (5'->3') |
|---|---|---|
| *sma-2* | TCCACCAGGAGTTCCAACAT | ACCTGTTCTCCGACTCTTGT |
| *sma-3* | GAGAACACACGGATGCATATTGG | ACTGTGCGGTGGTATTCGG |
| *sma-4* | GATGCTCCGACGTTCTCGAT | CGCATCCTGTCAACTCCACT |
| *act-1* | GCCGGAATCCACGAGACTTC | TCTGGTGGGGCGATGATCTT |

Supplementary Table 3. Survival assay summary

| Bacteria | Nematode strain | Trial 1 | | | | Trial 2 | | | | Trial 3 | | | |
|---|---|---|---|---|---|---|---|---|---|---|---|---|---|
| | | test mean lifespan ± SE | n | control mean lifespan ± SE | n | test mean lifespan ± SE | n | control mean lifespan ± SE | n | test mean lifespan ± SE | n | control mean lifespan ± SE | n |
| *E. cloacae* | WT | 16.22±0.39 | 104 | 12.3±0.37 | 110 | 15.58±0.30 | 105 | 14.98±0.35 | 102 | 16.68±0.52 | 71 | 13.84±0.40 | 75 |
| | *dbl-1(-)* | 11.64±0.33 * | 105 | 11.87±0.38 | 111 | 13.67±0.41 ns | 112 | 14.36±0.36 | 104 | 11.66±0.47 * | 58 | 14.17±0.40 | 58 |
| *K. oxytoca* | WT | 14.34±0.31 | 120 | 12.3±0.37 | 110 | 16.16±0.27 | 110 | 14.98±0.35 | 102 | 16.41±0.30 | 75 | 13.84±0.40 | 75 |
| | *dbl-1(-)* | 12.55±0.32 * | 108 | 11.87±0.38 | 111 | 14.87±0.31 ns | 100 | 14.36±0.36 | 104 | 9.31±0.55 * | 62 | 14.17±0.40 | 58 |
| *S. marcescens* | WT | 15.09±0.28 | 107 | 15.19±0.23 | 111 | 12.40±0.38 | 115 | 12.3±0.37 | 110 | 13.43±0.39 | 113 | 14.98±0.35 | 102 |
| | *dbl-1(-)* | 8.56±0.25 * | 111 | 14.58±0.22 | 97 | 6.58±0.29 * | 108 | 11.87±0.38 | 111 | 6.21±0.30 * | 105 | 14.36±0.36 | 104 |
| *B. megaterium* | WT | 16.60±0.21 | 101 | 13.44±0.21 | 115 | 16.54±0.21 | 84 | 14.06±0.24 | 95 | 12.86±0.46 | 63 | 9.07±0.55 | 71 |
| | *dbl-1(-)* | 13.82±0.19 * | 119 | 14.2±0.18 | 116 | 15.25±0.22 * | 84 | 13.99±0.22 | 84 | 13.47±0.27 ns | 73 | 8.61±0.37 | 77 |
| *E. faecalis* | WT | 17.77±0.28 | 121 | 13.82±0.25 | 121 | 16.54±0.29 | 119 | 13.20±0.22 | 123 | 17.46±0.32 | 129 | 13.32±0.24 | 121 |
| | *dbl-1(-)* | 15.83±0.34 * | 75 | 13.20±0.29 | 99 | 13.25±0.39 * | 142 | 13.72±0.28 | 129 | 14.97±0.33 * | 126 | 13.58±0.23 | 122 |
| *S. epidermidis* | WT | 17.92±0.21 | 122 | 13.82±0.25 | 121 | 15.97±0.34 | 104 | 13.20±0.22 | 123 | 16.60±0.20 | 115 | 13.32±0.24 | 121 |
| | *dbl-1(-)* | 8.84±0.42 * | 138 | 13.20±0.29 | 99 | 11.88±0.35 * | 138 | 13.72±0.28 | 129 | 13.32±0.39 * | 143 | 13.58±0.23 | 122 |

*, log rank test $p$ <0.0001, ns, log rank test $p$ >0.01, comparing mean *dbl-1(-)* lifespan with wild-type lifespan on each bacteria

Supplementary Table 4. RAD-SMAD reporter activity in response to Gram-positive bacteria

| Bacteria | Nematode strain | % animals with no detectable fluorescence | | |
|---|---|---|---|---|
| | | Trial 1 | Trial 2 | Trial 3 |
| *B. megaterium* | WT | 100% | 100% | 100% |
| | *dbl-1(-)* | 100% | 100% | 100% |
| *E. faecalis* | WT | 95% | 22% | 21% |
| | *dbl-1(-)* | 62% | 40% | 65% |
| *S. epidermidis* | WT | 95% | 95% | 91% |
| | *dbl-1(-)* | 100% | 95% | 78% |

Supplementary Data 1. Gene enrichment analysis by WormCat of differentially expressed genes between wild-type and *dbl-1(-)* populations exposed to a) *S. marcescens* and b) *E. faecalis*

a) http://www.wormcat.com/static/dynamic/RGS_Mar-25-2021-05_56_38/sunburst.html

b) http://www.wormcat.com/static/dynamic/RGS_Mar-25-2021-05_58_42/sunburst.html

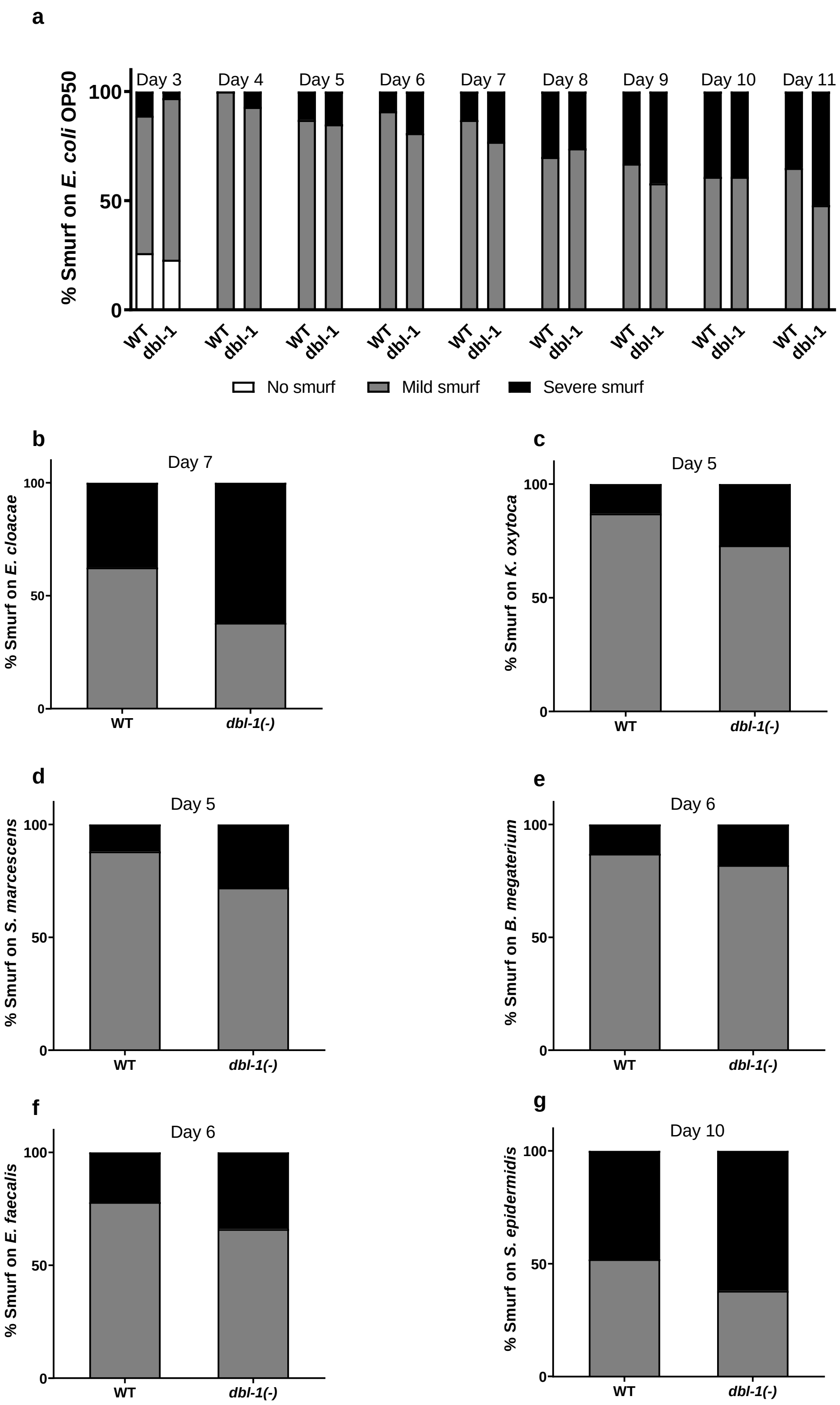
a
% Smurf on E. coli OP50
Day 3
Day 4
Day 5
Day 6
Day 7
Day 8
Day 9
Day 10
Day 11
100
50
0
WT
dbl-1
No smurf
Mild smurf
Severe smurf
b
Day 7
% Smurf on E. cloacae
WT
dbl-1(-)
c
Day 5
% Smurf on K. oxytoca
d
Day 5
% Smurf on S. marcescens
e
Day 6
% Smurf on B. megaterium
f
Day 6
% Smurf on E. faecalis
g
Day 10
% Smurf on S. epidermidis

Supplementary Figure 1. Loss of DBL-1 does not affect intestinal integrity upon exposure to specific bacteria. Wild-type and *dbl-1(-)* animals at the L4 stage were exposed to the following bacteria a) *E. coli* OP50 (control), b) *E. cloacae*, c) *K. oxytoca*, d) *S. marcescens*, e) *B. megaterium*, f) *E. faecalis*, or g) *S. epidermidis*. Intestinal barrier function was assessed using erioglaucine disodium salt a) over time or b-g) when *dbl-1(-)* populations neared their half lifespan. The leakiness of the intestine was assessed and scored as ‘1’ for no leakage/no Smurf, ‘2’ for mild leakage/mild Smurf, and ‘3’ for severe leakage/severe Smurf phenotypes. The fraction of animals indicating these phenotypes was calculated. One representative trial of at least three is presented. n= at least 10 per condition.

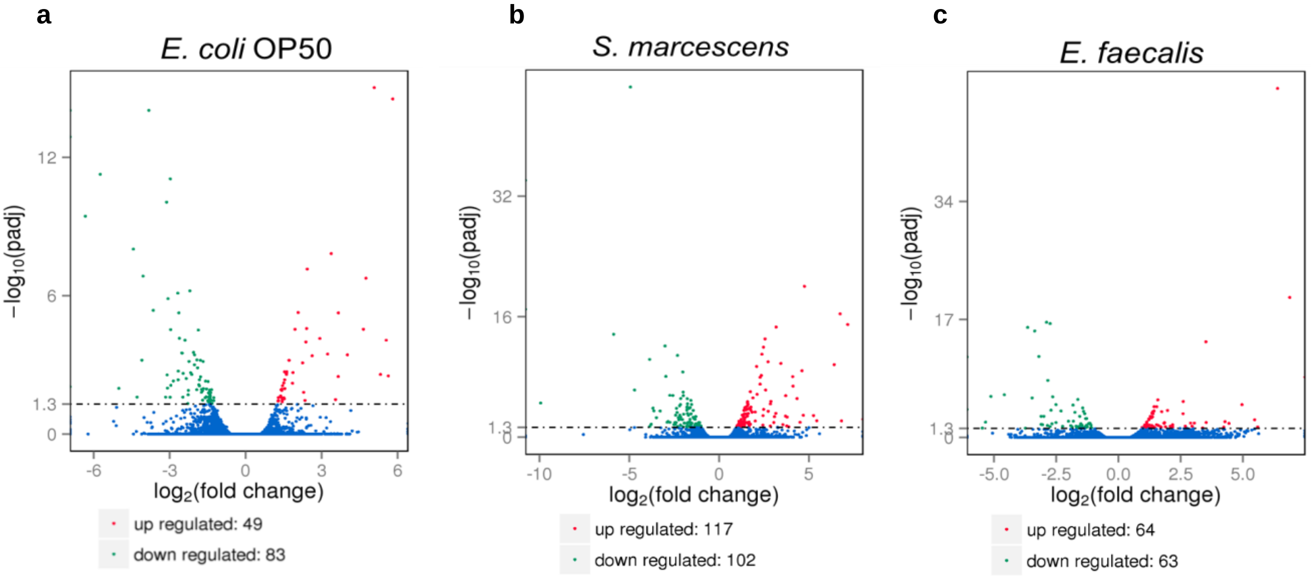


Supplementary Figure 2. DBL-1 regulates differential gene expression in response to Gram-negative and Gram-positive bacteria. Wild-type and *dbl-1(-)* animals were exposed to *E. coli* OP50 (control), *S. marcescens*, or *E. faecalis* at the L4 stage for two days. RNA-seq analysis volcano plots show differential gene expression in animals lacking DBL-1 exposed to a) *E. coli* OP50, b) *S. marcescens*, and c) *E. faecalis* in comparison to wild-type animals exposed to the same bacteria (adjusted *p*-value <0.01). Genes down-regulated in *dbl-1(-)* animals are represented in green, genes up-regulated in *dbl-1(-)* animals are represented in red, and genes with no change in expression are represented in blue.
